# Inferences from a network to a subnetwork and vice versa under an assumption of symmetry

**DOI:** 10.1101/034199

**Authors:** P.G.L. Porta Mana, E. Torre, V. Rostami

## Abstract

This note summarizes some mathematical relations between the probability distributions for the states of a network of binary units and a subnetwork thereof, under an assumption of symmetry. These relations are standard results of probability theory, but seem to be rarely used in neuroscience. Some of their consequences for inferences between network and subnetwork, especially in connection with the maximum-entropy principle, are briefly discussed. The meanings and applicability of the assumption of symmetry are also discussed.

PACS: 87.19.L-,87.19.lo,05.90.+m

MSC: 03B48,97K50

---

Remark: In this Note, whenever we use the term “probability”, we are not speaking about a frequency, or about some sort of (unmeasurable) physical property, but about a degree of reasonable belief, or plausibility, whose manipulation follows a well-defined and well-established logical calculus, “Bayesian theory” [1–27].

> **probable**, *a*. and *n*.
>
> **3. a**. Having an appearance of truth; that may in view of present evidence be reasonably expected to happen, or to prove true; likely.
>
> — Oxford English Dictionary [28]

> The correct probability is always that relative to the knowledge of the person who makes the probability-judgment.
>
> — C. D. Broad [29, ch. II, p. 150]

> When people say that the proposition “it is probable that p will occur” says something about the event p, they forget that the probability remains even when the event p does *not* occur.
>
> — L. Wittgenstein [4, p. 227]

## 1. Uncertainties about networks and subnetworks

If we are uncertain about the state of a network of neurons, what is our uncertainty about the state of a subnetwork? And vice versa, if we are uncertain about the state of a subnetwork, what is our uncertainty about the state of the whole network?

If our uncertainties are expressed by two probability distributions for the states of network and subnetwork, and these distributions satisfy a particular symmetry property, then they are related by precise and relatively simple mathematical formulae. These formulae are of essential importance when we want to make inferences about the whole network given data about the subnetwork and vice versa.

While well-known in survey sampling and in the pedagogic problem of “drawing from an urn without replacement”, such formulae are somewhat hard to find explicitly written in the neuroscientific literature. We have therefore decided to summarize them in this Note, as a reference, for the case of neurons modelled as “units” with binary {0, 1} = {inactive, active} states. The formulae also apply to any network or population of any kind of units modelled in the same way: e.g., the values of the physical connectivities of a neuronal network.

The formulae and the assumptions underlying them are given in the next two sections, followed by examples of their use in both inferential directions: network to subnetwork and reverse. We also discuss the meanings of the symmetry property, and conclude with a brief summary. Proofs of the formulae are sketched in an appendix.

Our notation follows ISO [30], ANSI/IEEE [31], and NIST [32] standards but for the use of the comma “,” to denote logical conjunction (“and”, “∧”), for the sake of typographic poise. Probability-calculus notation follows Jaynes’s book [22].

## 2. Formulae connecting network and subnetwork probabilities

### 2.1. Setup

Consider a network of *N* binary neurons with states (*X*_1_, …, *X_N_*) having fixed but unknown binary values (*R*_1_, …, *R_N_*), with *Ri* in {0,1}, vectorially written ***X*** = ***R***. For example, ***X*** can represent the state of a network at a particular time. We call the neurons “units” to lend some generality to our discussion. We shall make statements about the whole network of *N* units and about a subnetwork of *n* units; the word “network” will always refer to the *whole* network. The subnetwork states and their values are denoted by lowercase letters: (*x*_i_, …, *x_n_*) ≡ ***x*** and (*r*_1_, …, *r_n_*) = ***r***; but note that 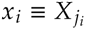; and 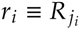 for some distinct *j*_1_, …, *j_n_*. We shall also make statements about the network-averaged state, or *network average:*

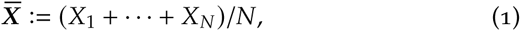

and the subnetwork-averaged state, or *subnetwork average:*

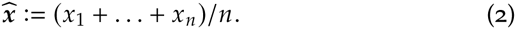

The quantities 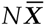 and 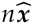 represent the total number of active units in the network and the subnetwork. Quantities like 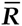 and 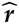 are defined analogously. The averaging operators 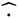 and 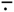 are also extended to averages of 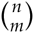 or 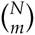 products of *m* states; e.g.,

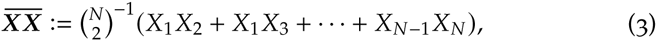

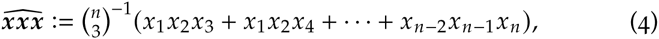

and so on.

### 2.2. Assumptions

Our uncertainty about the network state is represented by the joint probability distribution of the individual states, from which we can derive all other probabilities of interest. We denote it by

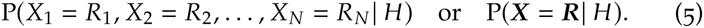

Such probability is conditional on our state of knowledge, i.e. the evidence and assumptions backing our probability assignments, denoted by the proposition *H*.

In the present discussion, *H* is a state of knowledge that leads to two specific properties in our probability assignments:

*H*_1_. *Permutation symmetry*, expressed as the invariance of the joint distribution (5) under arbitrary permutations of the units’s labels:

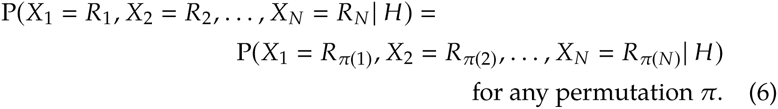

This property can reflect two very different states of knowledge: physical homogeneity of the network, or symmetry in our ignorance about the network. This property is called *finite exchangeability* in the Bayesian literature and its basis, consequences, and alternatives to it are discussed in § 4.

*H*_2_. The network average 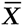 has a particular distribution *Q*:

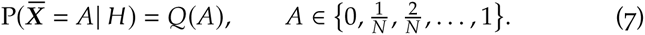

For the moment we are not concerned about the specific form of *Q* and about how it was assigned: it could, e.g., arise from maximum-entropy arguments [e.g.: 33–42] used with data on the network.

### 2.3. Formulae

The state of knowledge *H* has the following six (not independent) main consequences for our probability assignments:

***I***. Probability for the network state:

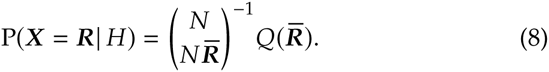

***II***. Probability for the state ***x*** of any subnetwork of *n* units:

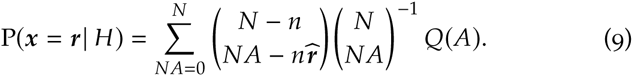

Note that the only summands contributing to this sum are those for which 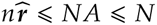; the others are zero because by definition 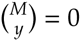 if *y* < 0. This remark applies to all the sums of this kind in the rest of this Note.

***III***. Probability for the subnetwork state conditional on a network state:

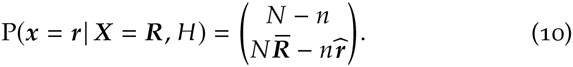

***IV***. Probability for the subnetwork average 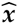:

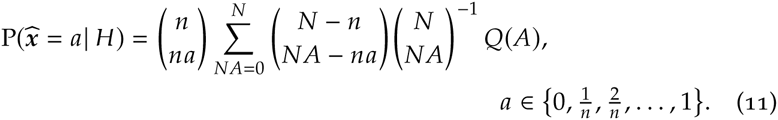

***V***. Probability for the subnetwork average conditional on the network average:

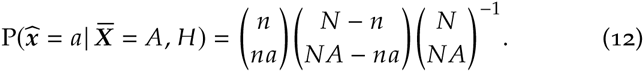

***VI***. The product of the states of any m distinct units from a given subnetwork,

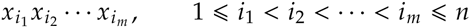

has an expectation equal to that of the subnetwork average of such products, is independent of the subnetwork size *n*:

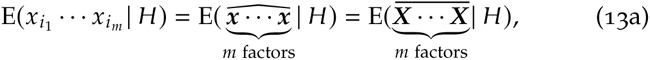

and has an explicit expression in terms of *Q*:

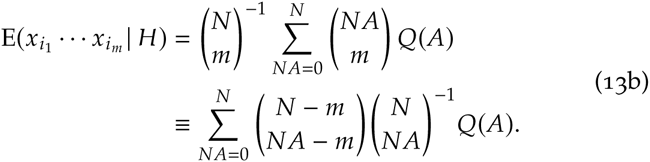

A useful relation connects the expectation of a product (13) and the *m*th *factorial moment* [43] of the probability distributions for the averages. The *m*th factorial moment of the subnetwork average 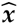 is defined by

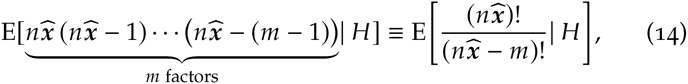

an analogous definition holding for 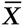. We have that

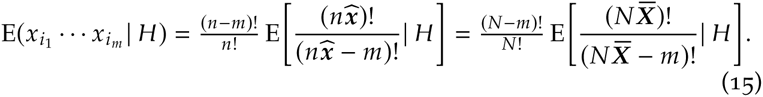

As a consequence of the above relation, the first three moments of the probability distributions 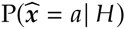 and 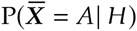, are related by

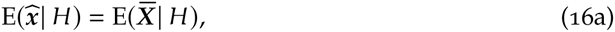

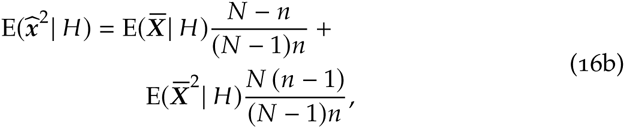

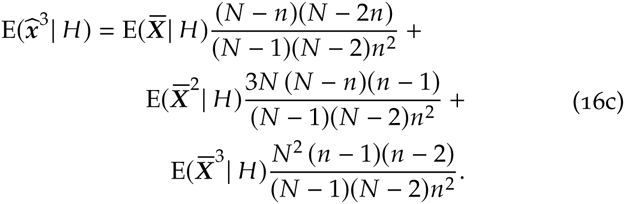

Relations for higher moments can be obtained recursively from eq. (15). In general, this means that the two sets of first *m* moments are related by a homogeneous linear transformation,

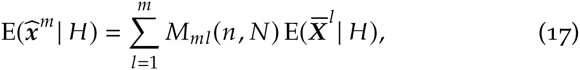

with a universal, lower-triangular transformation matrix *M_ml_*(*n*, *N*) that depends only on *n*, *N*, and the condition of symmetry (6).

As intuition suggests, we have

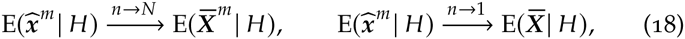

the latter because 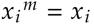, since states are {0, l}-valued.

The core of the six mathematical relations above are eqs (9) and (11). The latter expresses the probability for the subnetwork average as a mixture of hypergeometric distributions [22, ch. 3; 44, § 4.8.3; 45, § II.6], with parameters *N*, 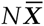, *n*, weighted by the probabilities 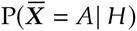 [cf. 46, § 4, esp. eq. (22)]. The connection between this mixture representation and the condition of symmetry (6) is well-known in the Bayesian literature [46–51].

Proofs of the above formulae are sketched in appendix A.

### 2.4. Asymptotic approximations and generalizations

Recall the definition of the Shannon and Burg [52; 53] entropies for a binary probability distribution (*p*, 1 − *p*) with *p* in [0, 1]

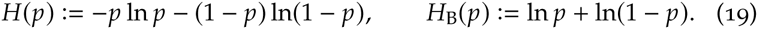

The binomial coefficient has an asymptotic form that involves the two entropies above [54–60]:

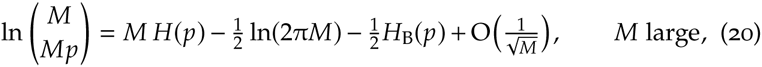

and this expression can be used to obtain asymptotic forms of the mathematical formulae (8)–(15) for large *N*, depending on the magnitudes of *n* and *n/N*. For example, if *N* and *n* are large and their ratio *k* := *n/N* finite, the probability distributions for the averages can be approximated (provided some regularity conditions) by continuum densities,

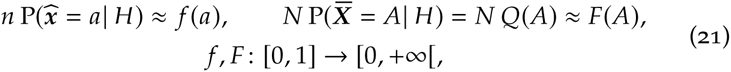

and sums by integrals, and eq. (9) takes on the approximate form

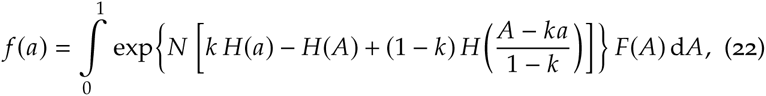

which shows an interplay of three entropies that depends on the ratio *n*/*N*. In some cases this allows us to approximate the integral by the value at a mode determined by a generalized entropy principle. (In fact, the standard maximum-entropy procedure can be derived from the probability calculus via a similar approximation [61].)

The relations above can also be generalized to *K*-valued states *X_i_* in {0, …, *K* − 1}, leading to the appearance of the generalized hypergeometric distribution [22, ch. 3; 44, § 4.8.3; 45, § II.6], or to real-valued states *X_i_* in **R**.

We do not pursue any of these approximations or generalizations in this Note.

## 3. Examples of inferential use of the formulae

### 3.1. From network to subnetwork

Let us illustrate with an example how the probability distribution for the subnetwork average 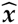, determined by eq. (11), changes with the subnetwork size *n*. Choose a network-average distribution 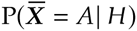 belonging to the exponential family [62, § 4.5.3; see also 63]:

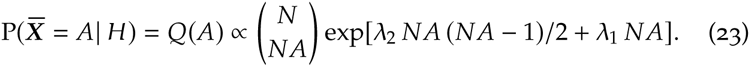

This is the form obtained from the principle of maximum relative entropy [e.g.: 33–42] with first and second moments as constraints and the reference distribution *Q*_0_ defined by 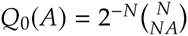, corresponding to a uniform probability distribution for the network state ***X***.

The probability distribution of eq. (23) is plotted in fig. 1, together with the resulting subnetwork-average distributions 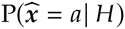, for the case in which *N* = 1000 units, λ_1_ = −2.55, λ_2_ = 0.005, and *n* = 10, 50, 100, 250. The distributions become broader as *n* decreases, and the minimum of the original distribution disappears; at the same time the finite-difference

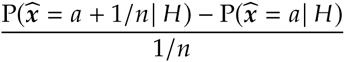

presents a sharp jump at this minimum when *n* ≈ 100.

**Figure 1.**
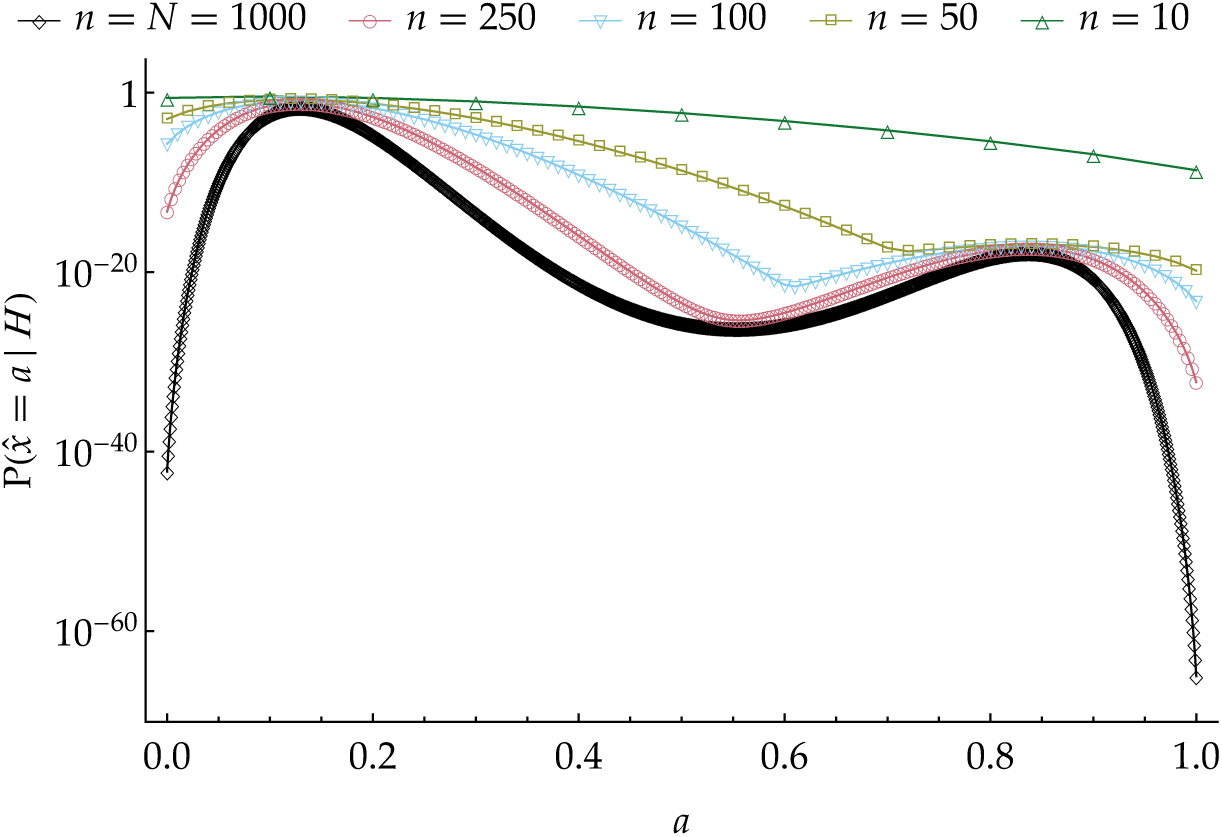
Probability distributions 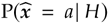 for different subnetwork sizes *n*, obtained from a network probability distribution 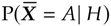 having the maximum-etropy form (23).

To the eye familiar with maximum-entropy distributions, the subnetwork-average distributions of fig. 1 do not look like maximum-entropy ones with second-moment constraints. In fact, they are not and *cannot* be:

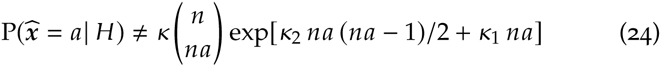

for any *κ*, *κ*_1_, *κ*_2_, unless *n* = 2. This impossibility holds more generally for any number of constraints *m* and subnetwork size *n* such that *m* < *n*. The reason is simple: suppose we have assigned a maximum-entropy distribution with *m* moment constraints as the distribution for the network average. If we want the same kind of distribution for a subnetwork of size *n*, we are free to play with *m* + 1 parameters (normalization included), but we must also satisfy the *n* + 1 equations corresponding to the marginalization (11). This is generally impossible unless 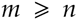. (Impossibilities of a similar kind appear in statistical mechanics, see e.g. ref. [64].)

This fact can be significant for recent works [e.g., 65–73] in which a maximum-entropy probability distribution with second- or third-moment constraints is assigned to relatively small subnetworks (*n* < 200) of neurons. If we assume that such subnetwork is part of a larger network, and assume the condition of symmetry (6), then the larger network *cannot* be assigned a maximum-entropy distribution with the same number of constraints. Vice versa, if we assign such a maximum-entropy distribution to the larger network, then none of its subnetwork of enough large size *n* can be assigned a similar maximum-entropy distribution. See ref. [74] for a broader discussion of this fact and of its consequences.

The dependence of the first four moments 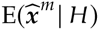 as a function of size *n* is shown in fig. 2. The moments become practically constant when *n* ≈ 100 or larger. The expectations of *m*-tuple products of states 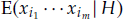, proportional to the factorial moments, are not shown as they do not depend on *n*.

**Figure 2.**
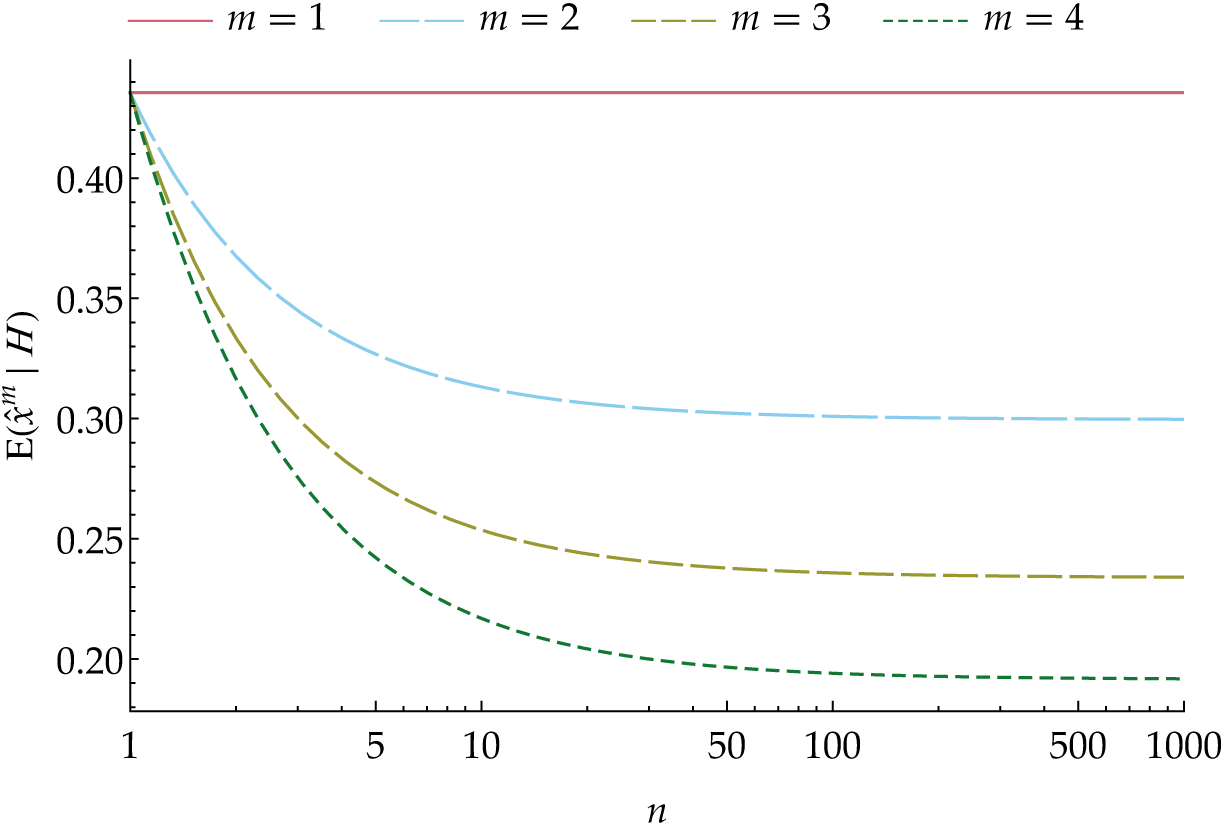
Moments of the probability distributions 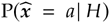 as functions of the subnetwork size *n*.

### 3.2. From subnetwork to network

We have seen that, given the condition of symmetry (6), the probability 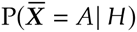 for the network average determines that of each subnetwork average, 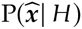, by the marginalization eq. (11). The reverse is trivially not true, since eq. (11), as a linear mapping from **R***^N^*^+1^ to **R***^n^*^+1^, with *N* larger than *n*, is onto but not into. Assigning a probability distribution 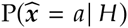 to a subnetwork average 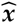 does not determine a network distribution 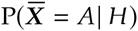: it only restricts the set of possible ones; this set can in principle be determined via linear-programming methods [23; 25; 75–77].

> Analogous situations appear in the truth-valued logical calculus: if the composite proposition A ⇒ *B* is assigned the truth-value “true”, then assigning *A* the value “true” also determines the value of *B*, whereas assigning *B* the value “true” leaves the value of *A* undetermined.

The same linear-programming methods show that any inference from subnetwork properties to network ones must necessarily start from some assumptions *I* that assign a probability distribution P(**X** = ***R***| *I*) for the network states. The approaches to this task and reformulations of it have become uncountable: they include exchangeable models, parametric and non-parametric models, hierarchical models, general linear models, models via sufficiency, maximum-entropy models, and whatnot [e.g.: 5; 22; 62; 78–86]. We now show two examples, based on a maximum-entropy approach, that to our knowledge have not yet been explored in the neuroscientific literature. For a concrete application see [87].

**First example: moment constraints for the network**. Consider a state of knowledge *H*′ leading to the following properties:

*H*′_1_. the expectations of the single and pair averages 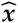 and 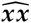 of a particular subnetwork have given values

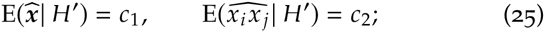

*H*′_2_. the network probability distribution P(***X*** = ***R***| *H*′) has maximum relative entropy with respect to the uniform one, given the constraints above.

Then the probability distribution for the network conditional on *H*′ is completely determined: it satisfies the symmetry property (6) and is defined by

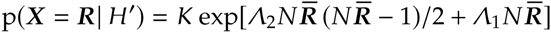

with *K*, *Λ_m_*, such that the distribution is normalized and

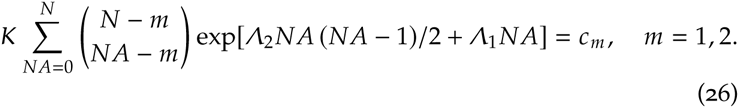

We omit the full proof of this statement: it is a standard application of the maximum-entropy procedure [e.g.: 33–36; 38–42], combined with the equality (13) of subnetwork and network expectations, e.g.

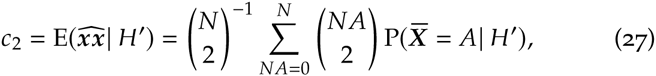

and with relations (8), (10). This example is easily generalized to any number *m* of constraints such that 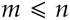.

Note again that, as remarked in § 3.1, the subnetwork from which the averages in the expectations (25) are calculated has a probability distribution 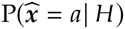 determined by the marginalization (11) and does *not* have a maximum-entropy form with the same number of constraints.

**Second example: subnetwork-distribution constraint**. Consider another state of knowledge *H*″ leading to the following properties:

*H*″_1_. the average 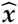 of a particular subnetwork has a probability distribution *q*:

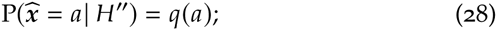

*H*″_2_. the probability distribution for the network, P(***X*** = ***R***| *H*″), has maximum relative entropy with respect to the uniform one, given the constraint above.

Then the probability distribution for the network given *H*″ is completely determined and satisfies the symmetry property (6):

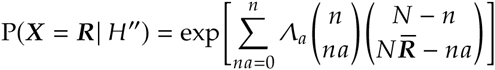

with *Λ_a_* such that

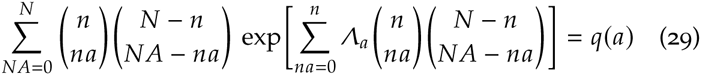

(the normalization constraint being unnecessary since *q* is normalized). This result is just another application of the maximum-entropy procedure with *n* + 1 (linear) constraints given by eq. (9), where the left-hand side is now given and equal to *q*(*a*).

This example is equivalent to the generalization of the previous one with *n* moment constraints, since knowledge of 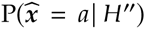 is equivalent to knowledge its first *n* moments.

The above examples do not mention how the values of the expectation constraints or of the subnetwork-average probability distribution can have been assigned. They cannot be assigned by a measurement, of course, since probabilities and expectations are not physical quantities and cannot be physically measured – they represent guesses of an observer and depend on the observer’s state of knowledge and assumptions. Rather, such values usually come from measurements made on “copies” – in a very general sense – of the states of the network; e.g., when we have a time sequence of them and measure the frequencies of their occurrence. Such situations can again be fully analysed with the probability calculus, and one can show [61] that the maximum-entropy formulae in the examples above are just limit forms of such an analysis, employing measured physical data *Δ* like, e.g., frequencies, and repeated applications of Bayes’s theorem,

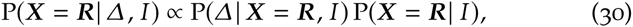

which updates the initial probability assignments on such data. But the discussion of this is again outside the scope of this Note.

## 4. On the symmetry property

The symmetry property (6) is called *finite exchangeability* in the Bayesian literature and, as was mentioned in § 2.3, its relation to the hypergeometric distribution in expression (11) for the subnetwork average is well-known [46–51].

This property can reflect two very different states of knowledge: either (a) knowledge that the network is somehow *physically homogeneous*, or (b) *complete ignorance* about the network’s homogeneity or inhomogeneity. In the second case we are saying that the indices or labels “*i*” of the units are *uninformative* – because, for example, we have no idea of how the units were labelled, hence we cannot presuppose any relation among the units, nor can we presuppose any structural or topological properties of the network they constitute. In the first case we are saying that the labels are *irrelevant*, even though they might be informative. The reason could be that, even if there is a connection between labels and, say, spatial locations of the units, each unit is nevertheless physically, homogeneously, identically connected to all the others; so we assume that spatial location does not play a relevant role.

An important consequence of the symmetry property is that no amount of new evidence about an *a*symmetry in the labelling of some units can lead to asymmetric predictions about the remaining ones. For example, if we have new data *Δ* saying that the first *n* units are in state 1 and the last *n* in state 0,

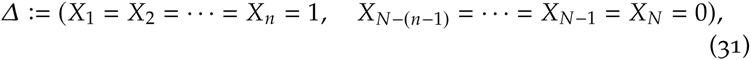

with *n* large, our updated probabily distribution will still assign (as can be shown using eqs (6) and (8)) the same probability for their respective neighbouring units with labels *n* + 1 and *N* − *n* to be in state 1 or 0:

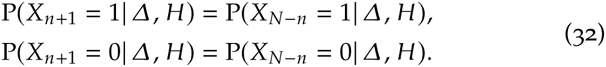

This may seem unreasonable – we would now say that the unit *n* + 1 is more likely to be in state 1, as the first *n* are, than in 0; and that the unit *N* − *n* is more likely in state 0, as the last *n* are, than in 1; i.e.

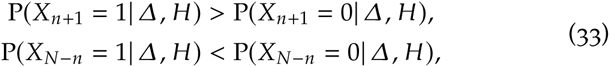

which is equivalent to 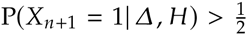, 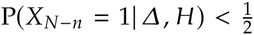; but the equalities (32) make this impossible.

The two very different motivations behind the symmetry property – lack of information and lack of relevance – can of course be handled differently by the probability calculus, in such a way that the appearance of asymmetry in new data leads to asymmetry in updated predictions. But this requires a more complex set of assumptions than those embodied in *H*; it requires, in particular, some sort of probability distribution for the degree of physical symmetry or homogeneity of the network, appropriately quantified.

The moral is that we should use the symmetric assumption *H* only if we can safely exclude the presence of inhomogeneity or are uninterested in detecting its presence. Otherwise, we must resort to more appropriate (and complex) assumptions. If we repeatedly observe new values that happen to have a very low probability according to the updated distribution P(***X*** = ***R***|*Δ*_1_, *Δ*_1_, …, *H*), this could be an indication that the symmetry property is unreasonable. Any strong departure of higher powers of measured averages from their expected behaviour given in eq. (16) and illustrated in fig. 2 can also be an indication that the symmetry property may have to be abandoned; hence the usefulness of eq. (16).

## 5. Summary and remarks

The main point of this Note was to explicitly collect and illustrate the mathematical formulae I–VI, § 2.3, between the probabilities assigned to the state of a network of neurons (or similar entities), and those assigned to the state of a subnetwork thereof. The formulae hold in the simple case of binary states and under an assumption of symmetry. Such relations can be found in several classical texts on probability and statistics; but we deemed it useful to restate them in a neuroscientific context, given their fundamental importance in relating a whole to its parts.

We have indeed seen that these formulae lead to straightforward but in some cases unexpected consequences: e.g., if a network is assigned a maximum-entropy probability distribution, then its subnetworks *cannot* be assigned a maximum-entropy distribution of the same form, and vice versa. The formulae also readily suggest new ways of making inferences or of formulating starting assumptions. Discussion of these points is left to forthcoming works [74; 87].

We also hope to have given the readers a feeling of the agility and simplicity with which the probability calculus (“Bayesian theory”) leads us from assumptions to consequences (albeit sometimes with non-simple mathematics), and from consequences to the assessment of how suitable our initial assumptions are, as shown with the assumption of symmetry.

## A. Sketched proofs

Variants of the following derivations and combinatorial considerations can be found e.g. in [88, chs I–IV; 45, ch. II; 22, ch. 3]; see also [89].

To derive the joint probability distribution (8) from that for the network average (7), consider that if the network total is 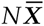, then 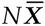 out of *N* units are active, and there are 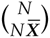 possible states for which this can be true; therefore

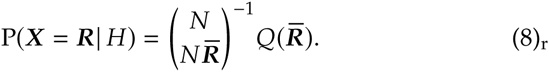

An analogous reasoning for *n* and 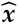 leads to an analogous equality,

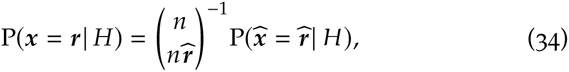

for the subnetwork.

Let us next consider the probability 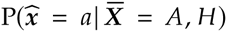 for the subnetwork average 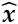 conditional on the network average 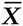. There are 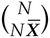 possible network states if the network average is 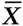, i.e. if 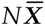 units are in state 1; the conditional probability of each is therefore 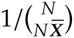, owing to the symmetry assumption (6). Now consider the subnetwork of the first *n* units. The conditional probability of having 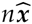 specific ones in state 1 is the sum of the probabilities of all states for which 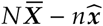 of the remaining *N* − *n* units are in state 1; there are 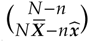 such states, all equally probable. Finally, there are 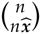 possible ways, all equally probable, in which 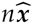 of the first *n* units can be in state 1. In formulae,

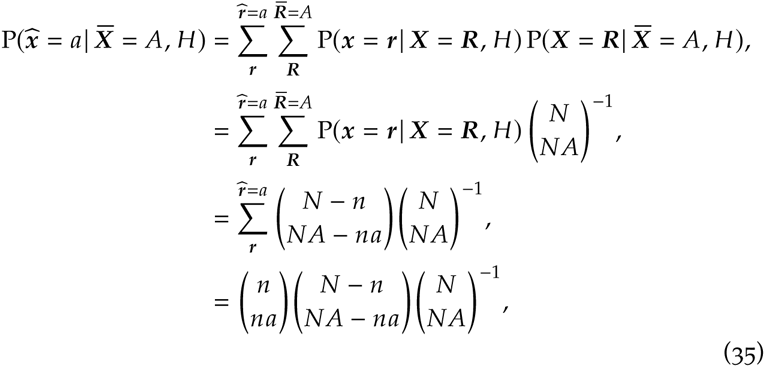

which is the conditional probability (12). Note that this is just a derivation of the hypergeometric distribution [22, ch. 3; 44, § 4.8.3; 45, § II.6], which describes the probability of, say, drawing a proportion of 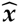 blue balls in *n* drawings without replacement from an urn with *N* balls, a fraction 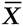 of which are blue.

The probability of a subnetwork average 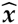 is then, by marginalization,

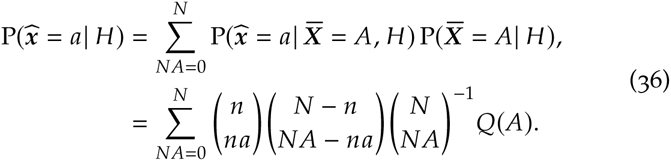

which proves the subnetwork-average formula (11). This formula, combined with eqs (34) and (8), leads to the conditional probability (10).

The independence of the expectation of products of states from the subnetwork size is trivial by marginalization:

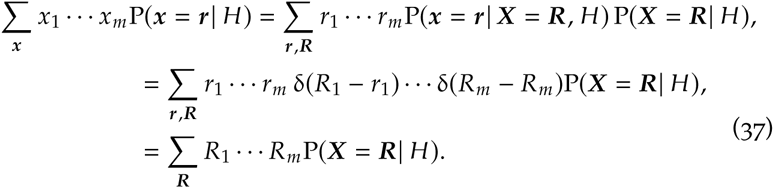

All such *m*-fold products have the same expectation by symmetry, therefore their subnetwork average will do, too, being an average of equal terms.

Now consider the sum of all distinct products of states of two units in the subnetwork:

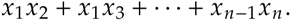

The terms in this sum are either 0 or 1. The non-vanishing ones are those with index pairs chosen from the 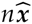 units of the subnetwork which are in state 1, and there are 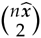 such choices, so the sum above is equal to 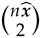. The sum has 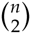 terms, so their average is 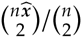. Generalizing the argument to products of *m* units, we have that

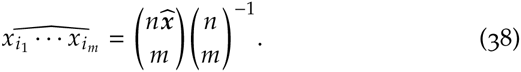

Then, using eq. (11),

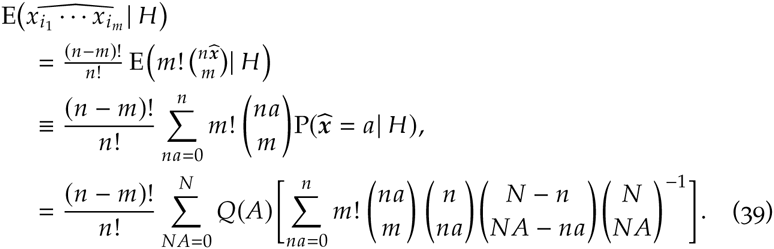

The expression in brackets is the *m*th factorial moment of the hypergeometric function, and is given by [43]

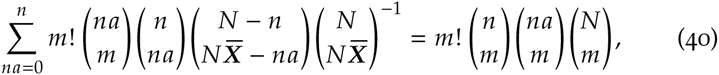

which combined with the previous equation yields the second line of eq. (13a); its last equality comes from the identity

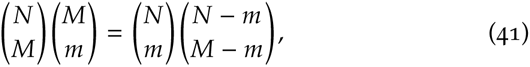

easily derived by writing the binomial coefficients in terms of factorials. Finally, eqs (16), relating the moments of the distributions for subnetwork and network averages, is obtained from the definition of moments,

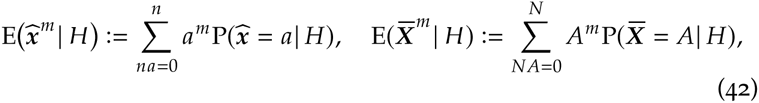

replaced in the equalities for the factorial moments (15), by recursively solving in terms of the moments of the network distribution.

## Acknowledgements

We thank Moritz Helias for many inspiring discussions, encouragement, and support; and gratefully acknowledge partial support by the Helmholtz young investigator group VH-NG 1028, the Helmholtz Portfolio Supercomputing and Modeling for the Human Brain, and the EU Grant 604102 (Human Brain Project). PGLPM thanks Buster Keaton for filling life with awe, Miri & Mari for continuous encouragement and support, the Forschungszentrum librarians for their prompt support in acquiring references, and the developers and maintainers of 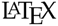, Emacs, 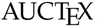, 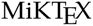, arXiv (except for its new “endorsement” system), Inkscape, Sci-Hub for making a free and unfiltered scientific exchange possible.

